# Adult male Lar gibbon sings the female great-call: A case study of inter-sex song production in a non-human primate

**DOI:** 10.1101/2024.08.21.608913

**Authors:** Judith Varkevisser, Stijn Berger, Judith van der Loo, Buddhamas Pralle Kriengwatana, Michelle Spierings

## Abstract

Gibbons are known as one of the most vocal non-human primates. They vocally advertise and reinforce their pair-bonds by singing complex duets, in which both the male and female perform a predetermined sex-specific set of song phrases, including the loud and elongated great-calls. Only females and sub-adult male gibbons have been previously observed performing great-calls. Once a male gibbon matures, he normally stops great-call production completely but continues singing other and less high-pitched song phrases. This case study describes a fully adult, castrated male lar gibbon (*Hylobates lar*, 32 years old, zoo-housed) who performs both male phrases of the duet, including the female great-call. The male regularly produced great-calls despite being in a, relatively weak, pair bond with a female conspecific. His great-calls adhered to the general structure of typical female great-calls but were shorter and had a lower maximum frequency. Notably, he produced these great-calls predominantly when the female was absent, especially when she was in their inside enclosure whilst he was outside. Behavioural observations indicate that the occurrence of great-calls by the male cannot be predicted based on pre-song behaviour or his interaction with the female. The recurrence of sub-adult singing behaviour in a fully grown adult is most likely due to his reduced testosterone levels. This study sheds light on the intricate dynamics of duetting and the unique occurrence of cross-sex song production in gibbons, emphasizing the complexity of pair-bond communication in this species.

## Introduction

Vocal communication in animals is often determined by sex-specific differences in terms of repertoire (Fischer & Hammerschmidt, 2020) and acoustic structure (Ey et al. 2007a; Heymann, 2011; Masataka & Fujita, 1989; Norcross & Newman, 1993; Norcross et al., 1999; Patel & Owren, 2007; Seyfarth et al., 1980). In some cases there is overlap in repertoire between males and females, but differences in anatomical features involved in sound production cause the structure of the vocalisations to be a reliable indicator of the sex of the producer (Ey et al., 2007b; Fitch, 2000). In species where males and females are less physiologically distinct, it is most commonly the repertoire that is determinedly sex-specific, and pair-bonded individuals often engage in antiphonal vocalization (Geissmann & Nijman, 2006; Marler & Tenaza, 1977; Marshall et al., 1972).

This antiphonal structure, or turn-taking behaviour, is usually temporarily coordinated (Pika et al., 2018; Ravignani et al., 2019; de Reus et al., 2000) and creates a back-and-forth communication system between two individuals. Antiphonal communication occurs in a diverse range of taxa, including birds (Thorpe et al. 1972; Hall, 2004), insects (Bailey, 2003), frogs (Tobias et al., 1998) and many nonhuman primates (tarsiers; Burton & Nietsch, 2010; titi monkeys; Müller & Anzenberger, 2002; gibbons and siamangs; Geissmann & Orgeldinger, 2000; Geissmann, 1999, 2002; Keith et al., 2009). The possible functions of song, a form of antiphonal communication, are all of a communicative nature. Song contributes significantly to territorial advertisement and mate attraction (Geissmann, 1999; Geissmann & Orgeldinger, 2000). A secondary function of these vocal alternations might be to maintain social bonds, both within the pair as within the family group, something that is seen specifically in gibbons (Wachtmeister, 2001; Terleph et al., 2015; 2016).

Gibbons live in the tropical forests in Southeast Asia, forming monogamous family groups that are territorial in nature (Chivers, 1976; Geissmann, 1995; Marshall & Sugardjito, 1986; Cowlishaw, 1992). These family groups typically consist of one male, one female, and their offspring. Upon reaching sexual maturity, offspring are driven to find new territories and mates (Brockelman et al., 1998). Gibbons mostly produce their coordinated songs during the early morning hours, probably because the conditions for sound transmission and visibility are optimal at that time (Morton, 1975). Their song contains information about the caller’s physical condition and identity (Clink et al., 2017; Terleph et al., 2015) and is known for their complex, antiphonal and sex-specific vocal structure. For example, the structure of lar gibbon songs is characterized by elements like notes and phrases, with great-call sequences representing a distinct, rapid pitch-rising pattern in female songs (Geismann, 1984). Duet sequences typically feature introductory “Hoo” notes, a dominant female sequence, a male “sharp wow” note, female great-calls, interlude sequences, and male solos (Raemaekers et al., 1984).

Multiple studies suggest that gibbon songs are mainly genetically determined (Geissmann, 1984; Brockelman & Schilling, 1984). Despite this, there is ongoing uncertainty about their learning abilities, particularly regarding duet coordination (Terleph et al., 2016). In agile gibbons (*Hylobates agilis*), older female offspring that demonstrate a heightened level of social independence produce vocalizations that are more closely aligned in timing with their mothers’ calls compared to their younger counterparts (Koda et al., 2013). However, like most primates, gibbons have not been previously found to improvise or imitate novel phrases, although they exhibit flexibility in the timing and pitch of their vocalisations.

While gibbons typically produce sex-specific song, rare cases of individuals singing the sex-specific phrases of the opposite sex have been documented. For instance, social circumstances were observed to induce male songs in female gibbons (Geissmann, 1984). Additionally, juvenile and sub-adult males may temporarily sing female-specific great-calls before transitioning to male-specific songs (Hradec et al., 2021; Koda et al., 2014). However, there are no reports, thus far, of fully adult males singing the female-specific great-call.

Here we report the first recorded vocal repertoire and behaviour of an adult male lar gibbon which produces the female great-call. This individual was born in captivity and currently lives with one female gibbon. He was observed producing female great-calls after his castration, although there are no records of his vocalisations before. To better understand the flexibility of song production in primates, we describe the acoustic structure of the great-call and the great-call containing song bouts of the male and female individual and provide an overview of the context of when these great-calls are produced.

## Material and methods

### Subjects

The study involved two adult lar gibbons (*Hylobates lar*), one male and one female, living in an enclosure at Safaripark Beekse Bergen, in the Netherlands. The male, Ori, was born in captivity on July 30th, 1987, at ZOO Antwerpen in Belgium, and subsequently transferred to Safaripark Beekse Bergen on May 20th, 1993. He underwent castration on November 21st, 2013. The female, Muguai, was born in captivity on January 12th, 2001, at Wildlands Adventure Zoo Emmen in the Netherlands and was relocated to Safaripark Beekse Bergen on October 24th, 2019. The male previously formed a strong pair-bond with another female, which lasted for 24 years until her demise on March 6th, 2017. The female in this study, Muguai, had a prior pairing with another male until she moved to Beekse Bergen.

The enclosure housing Ori and Muguai featured both an indoor and an outdoor area spanning 500 m^2^ (figure 1). During the day the gibbons where allowed access to both areas. Several trees (max 8m tall) and ropes were available for climbing (figure 2). There were no other gibbons in the enclosure. The gibbon enclosure is positioned on an island without enclosures containing other species around it (figure 1). However, the enclosures for chimpanzees, crested mangabeys, gorillas, and lions are within audible distance.

**Figure 1.**
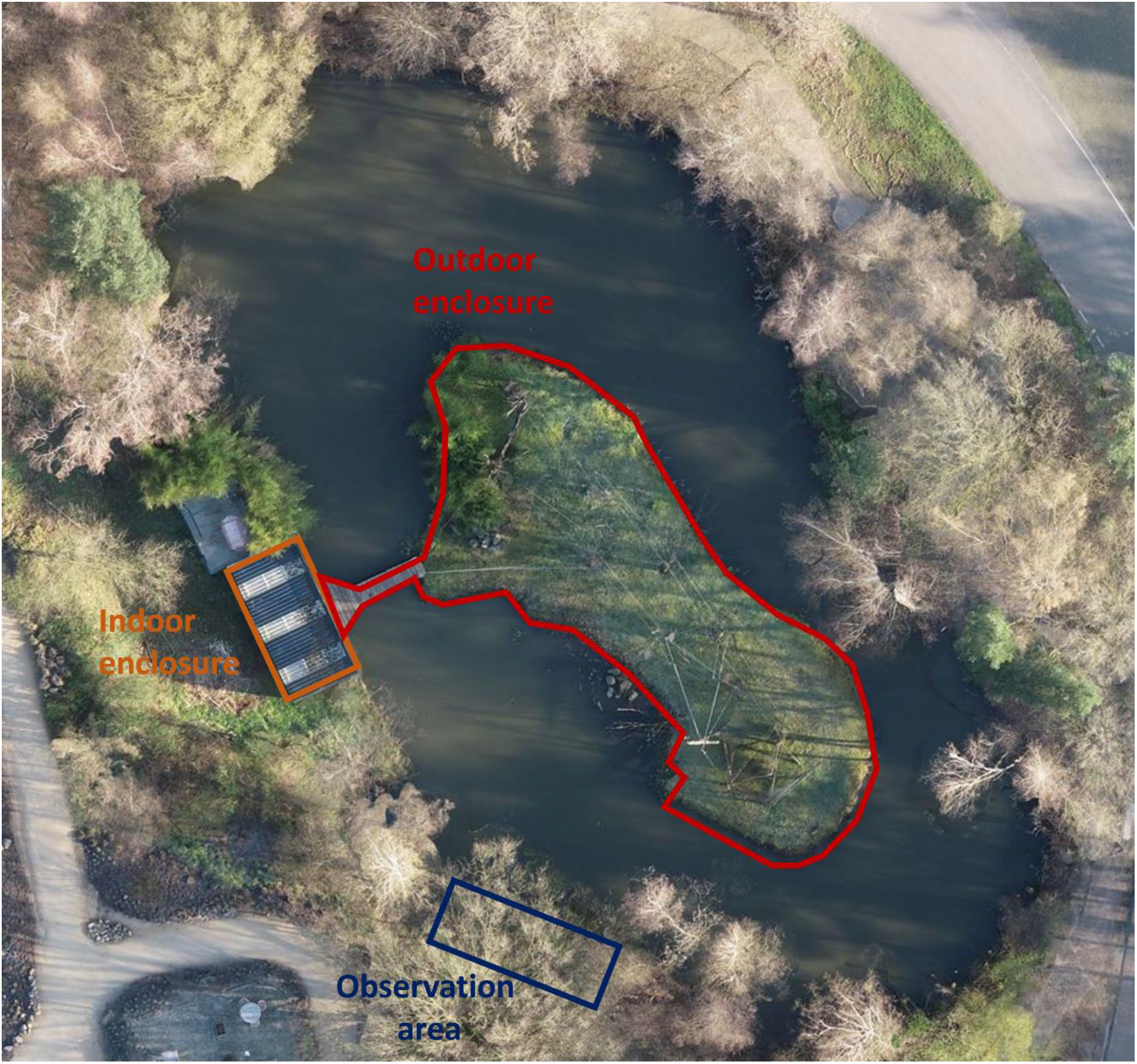
Top view of the indoor and outdoor areas of the gibbon enclosure in Beekse Bergen, and the area from which the audio recording and behavioural observations were made (‘Observation area’)

**Figure 2.**
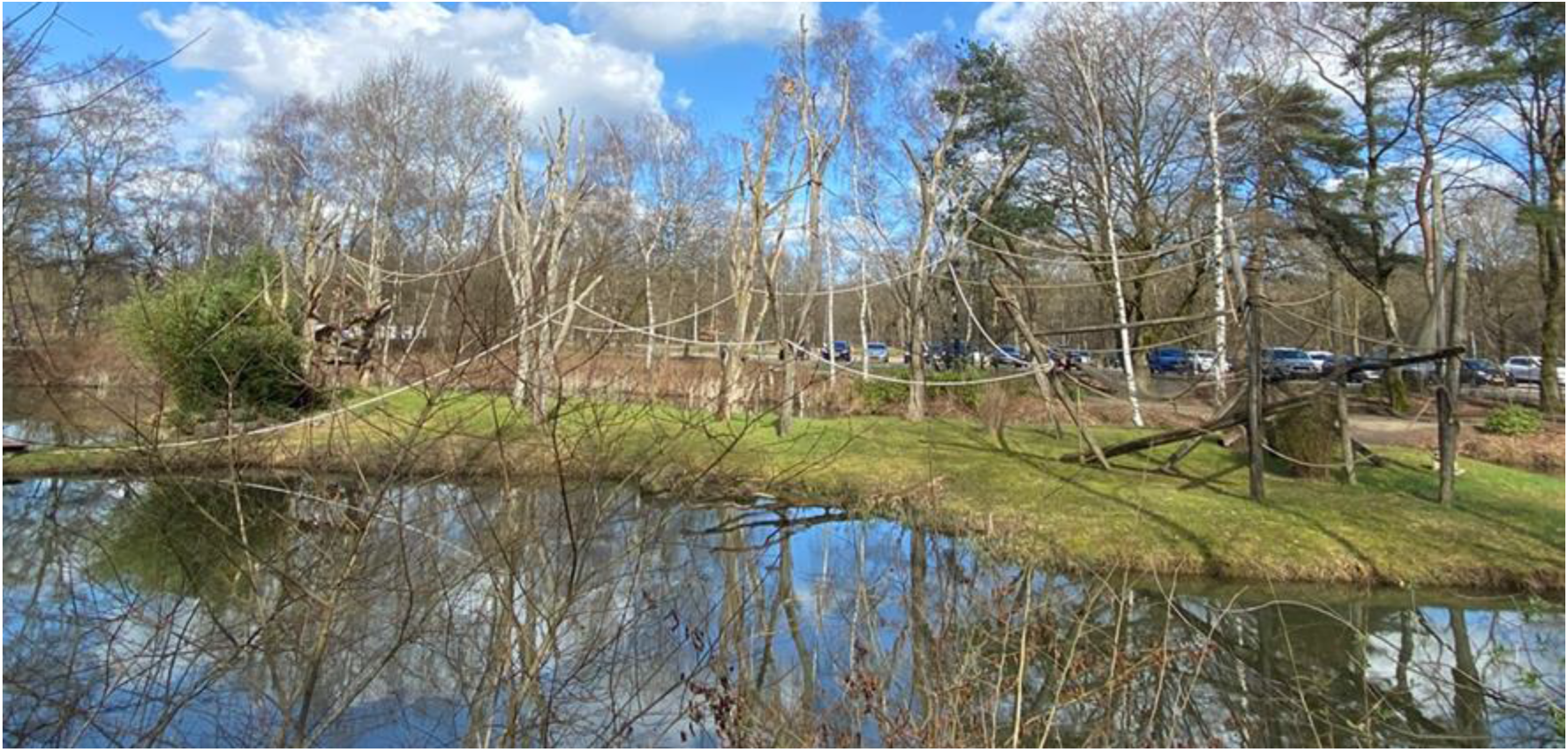
Picture of the outdoor area of the enclosure housing the male and female gibbon.

### Audio recordings

The gibbons were observed for a total of 29 days, within the period from March 28^th^ to May 12^th^ 2023. On the observation days, continuous audio recordings were made between approximately 07:00 and 14:00 o’clock (or until it started raining) with a unidirectional microphone (Sennheiser) with a Tascam DR-40 X Linear PCM recorder. The microphone was directed at the vocalising individual and notes were taken of which individual vocalized when. This ensured that the recorded songs could later be accurately attributed to either the male or female. The distance between the individuals and the microphone was between 25 and 35 meters, depending on the location of the gibbons. All recordings were cleaned with a high pass filter at 150Hz and were cut and annotated using Praat (Boersma & Weenink, 2008).

### Song Analysis

Praat (version 6.4.04) was used to create spectrograms of the recordings (settings: fast Fourier transformations with 1000 time and 250 frequency steps, 0.005s window length, dynamic range 50 dB, Gaussian window). Following Raemaekers et al. (1984), a note was defined as a continuous tonal sound produced by one individual and a bout as a period of calling by an individual separated from other periods of calling by at least ten minutes. In line with Terleph et al. (2015), the onset of a great-call was defined as the start of the first note that was more than 1 second long and the end of a great-call as the end of the first note following the climax note (defined as the note with the highest frequency excluding ‘high notes’) that was less than 700 ms long. For each great-call, the number of notes it contained was determined and its duration was established by examining the interval between the onset of the first and the end of the last note (figure 3). In Praat, spectral slices of the climax note of each great-call were created to determine the highest F0 frequency in these notes. For each bout containing a great-call, the duration of the calling preceding the onset of the first great-call of that bout was determined.

**Figure 3.**
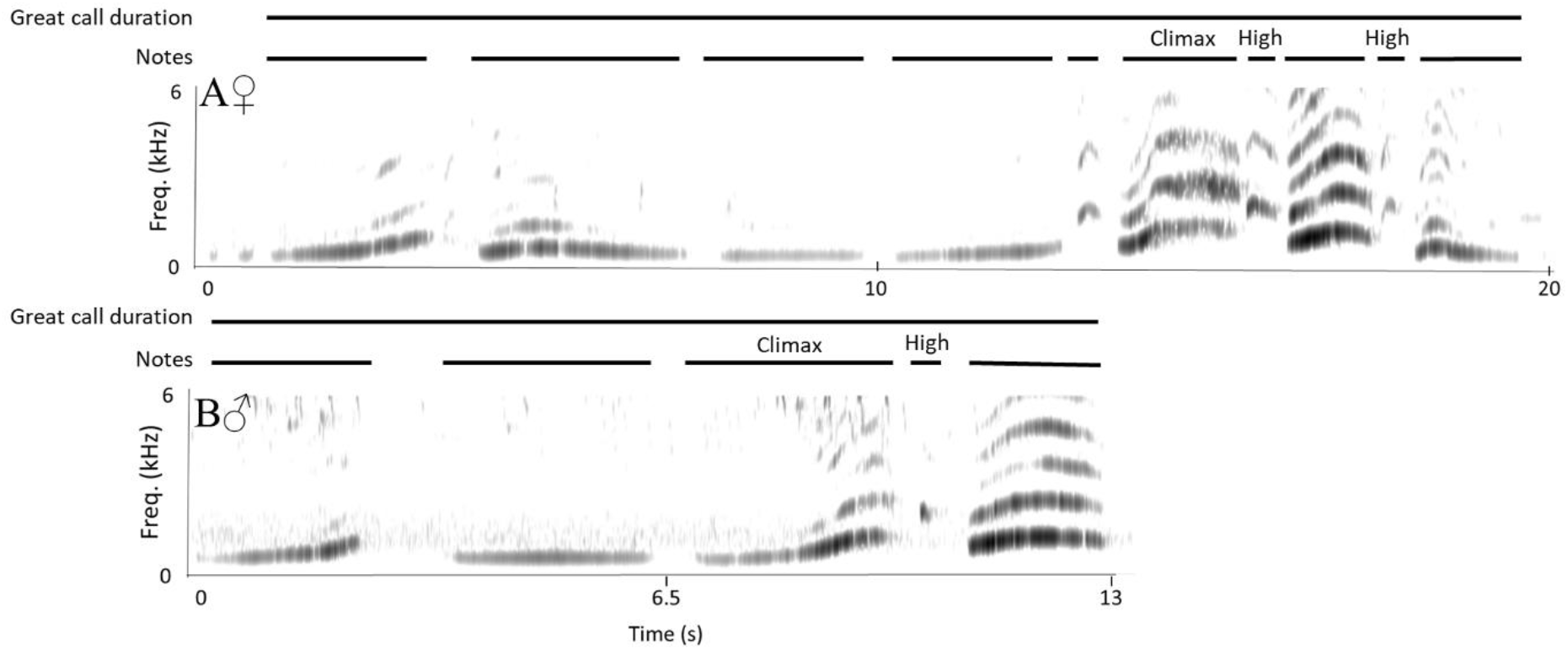
Spectrogram of a representative great-call produced by the female (A) and male (B) gibbon. The lines above the spectrograms indicate the individual notes in the great-call, the climax and high notes and how the great-call duration was measured.

### Behavioural analysis

Behavioural observations were conducted on the 29 observation days between 07:00 and 14:00 when the gibbons were in the outdoor area of their enclosure. These observations were documented through video (JVC camcorder GZ-R415G) recordings, as well as direct behavioural monitoring with ZooMonitor (version 4.3.9; Ross et al., 2016). To investigate the behavioural and social context in which the male individual produced bouts with great-calls, we analysed behavioural observations made 30 minutes before the male individual started singing, during its songs and 30 minutes after singing. During each observation period (before, during and after song), the social and solitary behaviours of both individuals (measured as social proximity and position in outdoor area) were recorded every 20 seconds. Social proximity data were categorized into five levels: 0 m (contact), 0 – 2 m (contact possible), 2 – 4 m (no contact possible), 4+ m (far away), and out of sight (one of the individuals is inside). Besides this, the position of the male individual in the outdoor area was categorized into five height levels: on the ground, low (just above the ground, below the ropes stretched in between the trees in the enclosure), middle (in between the ropes), high (above the ropes, in the top of the trees in the enclosure) or out of sight (inside). The duration of pre- and post-song periods on the one hand and the during song period on the other, were different, as the pre- and post-song periods were fixed at 30 minutes while the during song period was variable depending on the duration of the bout. To account for these different durations, the number of intervals during which a behaviour was observed within a period was divided by the total number of intervals in that period and multiplied by 100 to express the behaviour’s occurrence as a percentage.

## Results

### Song analysis

During the observational period, we recorded 18 bouts produced by the female individual, of which 11 bouts (61.1%) contained great-calls. These bouts contained 58 great-calls in total (an average of 5.3 great-calls per bout). We recorded 24 bouts from the male individual, with 17 bouts (70.8%) containing great-calls. These contained 40 great-calls in total (an average of 2.4 great-calls per bout).

Figure 3 shows spectrograms of representative examples of the great-calls produced by the female (A) and male (B) individual. The female’s great-call starts with long notes with small frequency modulations, followed by an alternation of loud notes with larger frequency modulations and high notes. The great-calls produced by the female contained between 8 and 11 notes (average ± SD = 9 ± 0.82 notes). The male’s great-calls also start with long notes with small frequency modulations followed by two loud notes with larger frequency modulations and one high note in between. All great-calls produced by the male contained these same 5 notes.

The female’s great-calls were on average (± SD) 18.44 (± 1.30) seconds long, while those produced by the male were on average (± SD) 12.10 (± 0.55) seconds long. The highest F0 in the climax note was on average (± SD) 1320 (± 61.79) hertz in the female’s great-calls and 1247.63 (± 37.11) hertz in the male’s great-calls. The first great-call in the bout occurred on average (± SD) after 149.69 (± 34.24) seconds in the female individual and after 481.09 (± 222.29) seconds in the male individual. Figure 4 shows an example of a bout containing two great-calls produced by the male.

**Figure 4.**
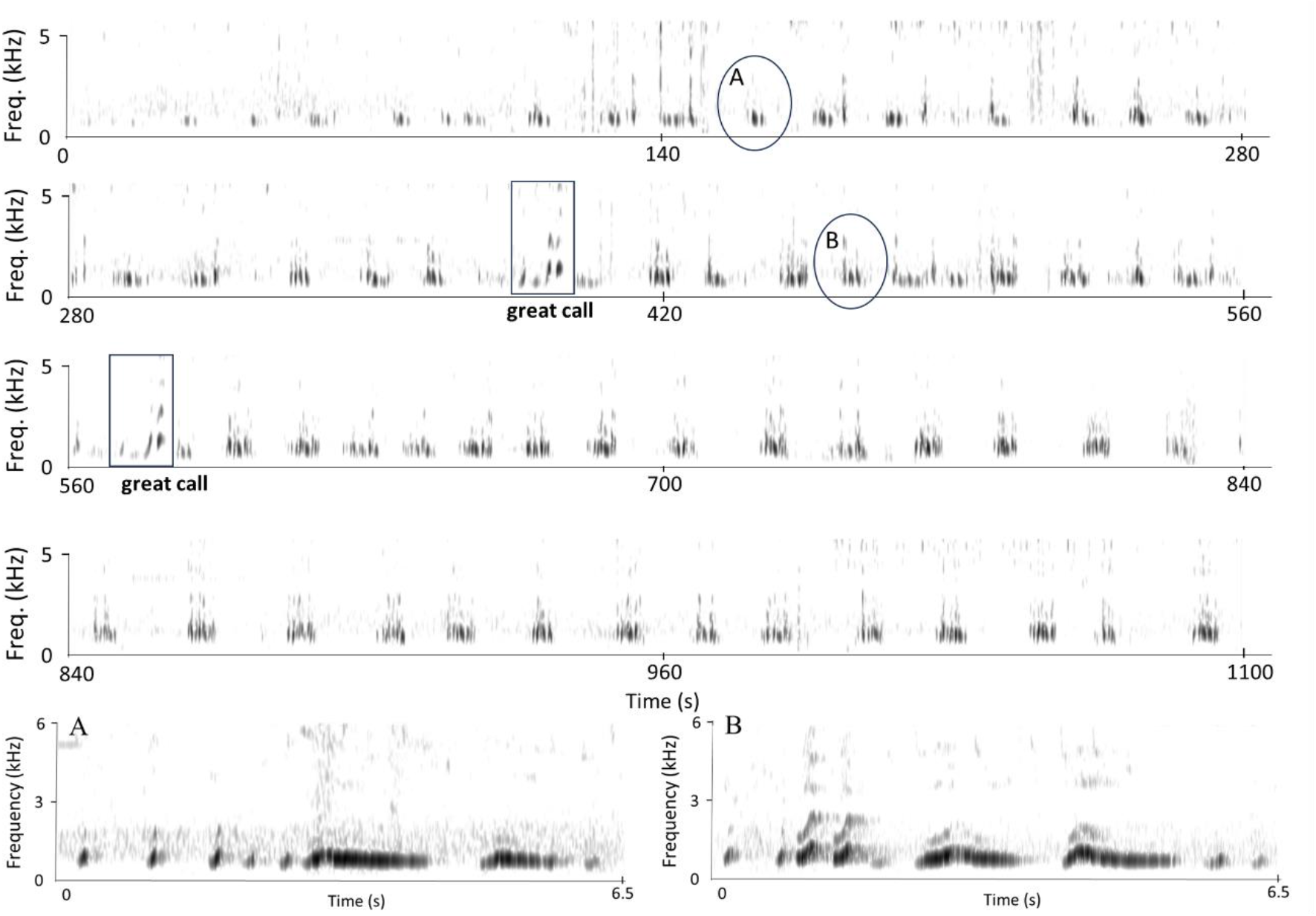
Spectrogram of a representative bout produced by the male gibbon. The bout contains two great-calls, which are indicated with a square. Two representative phrases from the part of the bout before the first great-call (A) and from the part in between the two great-calls (B) are shown in enlarged spectrograms (circles indicate their position in the bout).

Table 1 summarizes the differences between the great-calls and great-call containing bouts produced by the female and male individual. The average number of notes, duration and highest climax note F0 of the female’s great-calls fell within previously reported ranges for 14 wild female Lar gibbons calling individually (Terleph et al., 2015). The male’s great-call contained fewer notes, was shorter and had a lower maximum F0 compared to these previously reported female great-calls (Terleph et al., 2015).

**Table 1.** Characteristics of the great-calls and great-call containing bouts produced by the female and male individual.

|  | Female individual<br>(n = 11 bouts, 58 great-calls) | Male individual<br>(n = 17 bouts, 40 great-calls) |
| --- | --- | --- |
| | mean ± SD (range) | mean ± SD (range) |
| Number of notes in great-call | 9 ± 0.82 (8 – 11) | 5 ± 0 (5) |
| Duration of great-call (in s) | 18.44 ± 1.30 (16.24 – 21.11) | 12.10 ± 0.55 (10.72 – 13.15) |
| Highest F0 in climax note (in Hz) | 1320 ± 61.79 (1092 - 1463) | 1247.63 ± 37.11 (1202 - 1385) |
| Duration until first great-call<br>appeared during the bout (in s) | 149.69 ± 34.24 (94.30 –<br>202.51) | 481.09 ± 222.29 (72.62 –<br>867.41) |

### Behavioural analysis

The behaviour of both individuals indicate that the two gibbons did not have a strong pair bond. The individuals did not duet during any of the observations, nor was there allogrooming during any of the analysed observation intervals. Other affiliative behaviours, such as close proximity (being 0 to 2 m apart from each other) only occurred during 0.93% of these intervals.

During the observed intervals where the male individual produced bouts containing great-calls, the female individual was either out of sight (97.28% of the intervals, figure 5) or at more than 4 meter distance from the male (2.27%). In the half hour before and after these bouts, the female was less ‘out of sight’ (before: 56.47%, after: 62.59% of the intervals). The male individual mainly produced bouts containing great-calls while positioned in the treetops (‘high’, 83.21%, figure 6). Before and after singing, the male spent much less time in the treetops (‘high’ before: 3.40%, ‘high’ after: 10.94% of the intervals). None of the observed behaviours showed a clear difference before and after great-call production of the male gibbon. For instance, the two gibbons would not be in closer proximity after the male had produced a song with a great-call compared to before he had produced the song.

**Figure 5.**
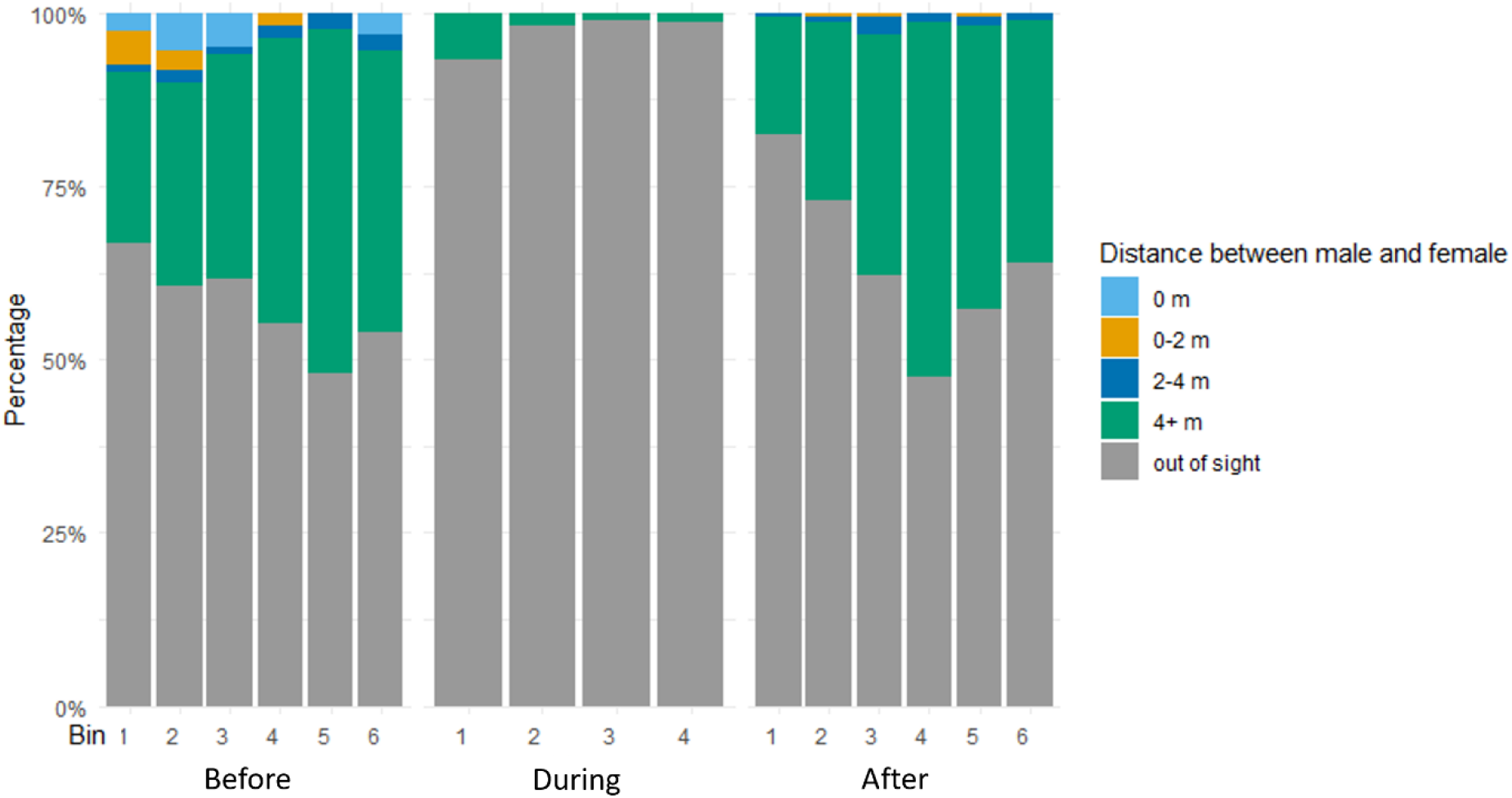
Percentage of observed intervals that the male (Ori) and female (Muguai) gibbon spent at different distances from each other half an hour before, during, and half an hour after the male’s great-call containing bouts. The half hour before and after song was divided into six bins of five minutes each, the period during singing was divided into four equally long bins (as the duration of this period varied from bout to bout).

**Figure 6.**
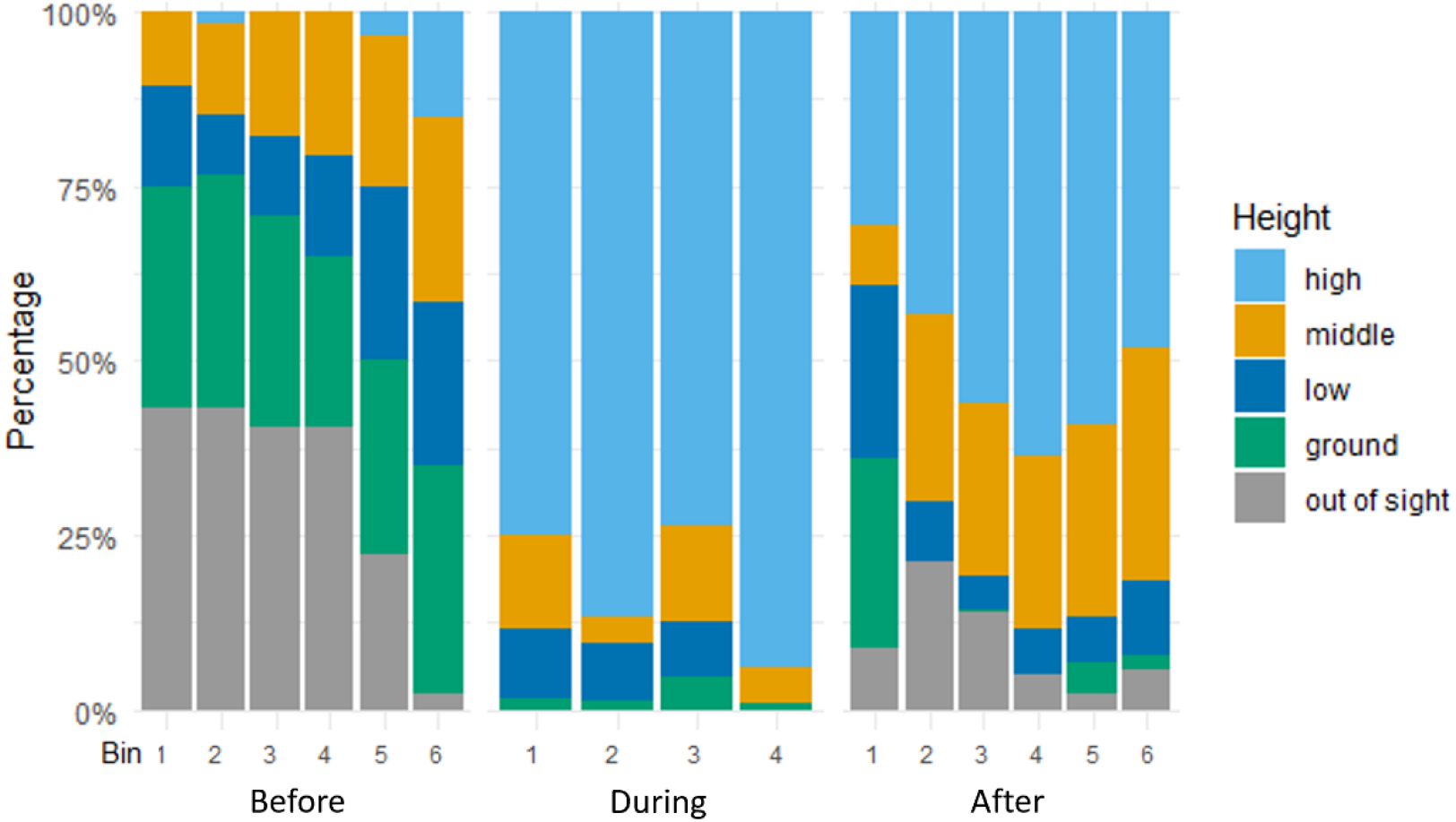
Percentage of observed intervals that the male spent at different heights in the enclosure half an hour before, during, and half an hour after producing great-call containing bouts. The half hour before and after song was divided into six bins of five minutes each, the period during singing was divided into four equally long bins (as the duration of this period varied from bout to bout).

## Discussion

The observation of an adult male lar gibbon exhibiting the female-specific great-call represents a unique and intriguing phenomenon within the context of gibbon vocal communication. The current findings contribute to the existing knowledge on sex-specific vocalizations in primates. The male in this study produced the female great-call, a behaviour not previously documented in fully mature males. This individual, born in captivity and castrated several years ago, challenges the traditional notions of gibbon vocalizations being strictly genetically determined. This means that under specific circumstances, previously produced vocalisations can resurface.

Reasons for the atypical singing behaviour of the adult male gibbon could hypothetically be related to 1) the desire to form stronger pair bonds with the female; 2) territorial defence; 3) absence of gonadal hormones. As antiphonal duetting plays a significant role in territorial advertisement, mate attraction and maintenance of social bonds within the family group (Wachtmeister, 2001; Terleph et al., 2015; 2016), it may have been possible that the male incorporated great-calls into his song to stimulate pair bonding. However, this hypothesis is not supported because the female was far away or out of sight when the male produced songs containing great-calls, and these songs did not seem to elicit a behavioural response in the female. Thus, singing songs with great-calls did not appear to increase social bonding behaviours between the male and the female. In fact, it might be possible that the male’s atypical songs hindered pair bond formation with the female (although not because she perceived that he was already pair-bonded and duetting with another individual, as gibbons identify individuals based on voice characteristics; Terleph et al., 2015).

The hypothesis that the male incorporated great-calls into his repertoire for more effective territory defence due to the weak pair bond with the female was also not supported because there were no other gibbons around. Rare cases of suspected interspecies interactions, such as vocalisations used for territorial defence against heterospecifics not normally considered as predators for that species, have also been documented (Domken et al., 2021). Future work could explore whether songs with the female great-call are more effective at repelling heterospecifics as well as the acoustical structural and temporal relationships between the male’s songs and vocalisations of other species in nearby enclosures, to ascertain whether the male is responding with songs to vocalisations or presence of other species.

We speculate that the atypical singing behaviour of this adult male was most likely linked to his castration. Recordings from the pre-castration period, while he was pair-bonded with his previous partner and from his sub-adult stage for direct comparison would have allowed us to be more confident in this conclusion. Nonetheless, our results generate interesting questions about the role of gonadal hormones in vocal repertoires and duetting. A wealth of evidence demonstrates the role of gonadal hormones in enabling and maintaining male song and stimulating male-typical songs in females (Gurney & Konishi, 1980; Vahaba et al., 2020; McLean et al., 2024). Hormone influence on song production plays a significant role in the context of duets. Steroid hormones, such as testosterone, have been shown to impact song production, as shown in white-browed sparrow weavers (Voigt & Leitner, 2013). However, very little knowledge exists for the role of gonadal hormones in suppression of call types in an animal’s vocal repertoire, and how its absence affects duetting behaviour.

In many species, juveniles and adults have a different vocal repertoire: juveniles often “lose” certain call types (e.g. begging calls) while “gaining” others (e.g. alarm calls), indicating that gonadal hormones could play a role in shifting from juvenile to adult repertoires. Gibbons, however, deviate from this pattern, as juvenile and sub-adult males temporarily sing female-specific great-calls before transitioning to male-specific songs (Hradec et al., 2021; Koda et al., 2014). While the conventional understanding has been that gibbon songs and duet coordination are primarily genetically determined (Terleph, 2016), age-related changes in temporal precision of vocalisations have been observed in female agile gibbons (Koda et al., 2013), Furthermore, social environments – potentially via short-term modulation of gonadal hormones – can cause female gibbons to produce male songs (Geissman, 1983). Thus, gibbons are an interesting model for understanding how genetic predispositions, endocrine function, and social environments influence vocal development and flexibility in production or usage.

In conclusion, the unexpected observation of an adult male lar gibbon producing the female great-call adds new insights to our understanding of gibbon vocal communication. This case challenges established notions of genetic determinants and highlights the need for further research into the interplay between genetic predispositions, life events, and social dynamics in shaping the vocal repertoire of vocal non-human primates.

## Statement of Ethics

This study was observational, and the observations only led to negligible interference in the daily lives of the gibbons.

## Acknowledgements

First of all, we would like to thank Prof. dr. Marco Gamba for discussing this project with us and providing us with several recordings of female gibbons singing great-calls. We further thank the students that worked on this project, Ella Baas and Bianca van der Burgh. We thank Beekse Bergen for allowing us to make recordings of their gibbons and prof. Thomas Geissmann for his input in the development of this project. This research was funded by FWF YIRG grant 10.55776/ZK66 & NWO Veni grant 212.264 (both awarded to MS).

